# Spatiotemporal Feature Selection Improves Prediction Accuracy of Multi-Voxel Pattern Classification

**DOI:** 10.1101/746735

**Authors:** Jeiran Choupan, Yaniv Gal, Pamela K. Douglas, Mark S. Cohen, David C. Reutens, Zhengyi Yang

**Affiliations:** Centre for Advanced Imaging, The University of Queensland, Brisbane, Australia; Queensland Brain Institute, The University of Queensland, Brisbane, Australia; Department of Psychology, USC Dornsife College of Letters, Arts and Sciences, University of Southern California, USA; Laboratory of Neuro Imaging, USC Stevens Neuroimaging and Informatics Institute, Keck School of Medicine, University of Southern California, Los Angeles, CA, USA; School of Information Technology and Electrical Engineering, The University of Queensland, Brisbane, Australia; Center for Cognitive Neuroscience, University of California, Los Angeles, California, USA; Neuropsychiatric Institute, University of California, Los Angeles, California, USA; Departments of Psychiatry and Behavioral Sciences, Neurology, Radiological Sciences, Biomedical Physics, Psychology, and Bioengineering, University of California, Los Angeles, California, USA.; California Nanosystems Institute UCLA School of Medicine, Los Angeles, California, USA; Brainnetome Center, Institute of Automation, Chinese Academy of Sciences, Beijing, China

**Keywords:** fMRI, Multi-Variate Pattern Classification, Spatiotemporal Feature Selection, Multi-band EPI, Random Forest, Support Vector Machine

## Abstract

The importance of spatiotemporal feature selection in fMRI decoding studies has not been studied exhaustively. Temporal embedding of features allows the incorporation of brain activity dynamics into multivariate pattern classification, and may provide enriched information about stimulus-specific response patterns and potentially improve prediction accuracy. This study investigates the possibility of enhancing the classification performance by exploring spatial and temporal (spatiotemporal) domain, to identify the optimum combination of the spatiotemporal features based on the classification performance. We investigated the importance of spatiotemporal feature selection using a slow event-related design adapted from the classic Haxby et al. (2001) study. Data were collected using a multiband fMRI sequence with temporal resolution of 0.568 seconds. A wide range of spatiotemporal observations was created as various combinations of spatiotemporal features. Using both random forest, and support vector machine, classifiers, prediction accuracies for these combinations were then compared with the single time-point spatial multivariate pattern approach that uses only a single temporal observation. The results showed that on average spatiotemporal feature selection improved prediction accuracy. Moreover, the random forest algorithm outperformed the support vector machine and benefitted from temporal information to a greater extent. As expected, the most influential temporal durations were found to be around the peak of the hemodynamic response function, a few seconds after the stimuli onset until ∼4 seconds after the peak of the hemodynamic response function. The superiority of spatiotemporal feature selection over single time-point spatial approaches invites future work to design systematic and optimal approaches to the incorporation of spatiotemporal dependencies into feature selection for decoding.

**Highlights:**

- Spatiotemporal feature selection effect on MVPC was assessed in slow event-related fMRI
- Spatiotemporal feature selection improved brain decoding accuracy
- From ∼2-11 seconds after stimuli onset were the most informative part of each trial
- Random forest outperformed support vector machines
- Random forest benefited more from temporal changes compared with support vector machine

## 1. Introduction

In conventional univariate functional Magnetic Resonance Imaging (fMRI) analysis, the objective is to find brain regions that show reproducible activation with the repetition of specific experimental conditions (Friston et al., 1994). In contrast, in Multi-Variate Pattern Classification (MVPC) approaches, the pattern of responses across multiple brain voxels that together carry information about different experimental conditions is sought (Haxby et al., 2001). In MVPC, the functional relationship between across-voxel patterns of activation and the experimental conditions is modeled using discriminative pattern recognition techniques; the experimental conditions are then predicted from the fMRI signal (see (Haynes, 2015) review of MVPC).

A potential advantage of the MVPC approach over classical univariate analysis methods is that a fixed HRF model does not need to be assumed or estimated. However, this benefit from MVPC is yet to be realized fully because event-related decoding studies generally extract features at fixed temporal delays, which are themselves determined based on a canonical HRF, following the stimulus onset (e.g. Douglas et al. (2011)).

Even in the context of block designs, it may be critically important to take into account fMRI temporal dynamics in addition to multivariate spatial information in MVPC. There is strong temporal correlation in the fMRI time series, especially due to the delay and smoothing from the HRF. Mourao-Miranda et al. (2007) studied the temporal dynamics for MVPC by training and testing a classifier using all temporally contiguous acquisitions in each block, effectively treating time as spatial information, to produce SpatioTemporal (ST) signals. They found a localized peak of response in the amygdala only at a specific time point in the block suggesting that temporal averaging of fMRI activity in a block (i.e. assuming that hemodynamic responses to the same stimulus are a *stationary process*) averaged out the effect of specific discriminating times in specific regions, and ignored the temporal profiles caused by the hemodynamic response.

One study investigated the effect of entire-trial ST temporal embedding on MVPC accuracy of slow event-related fMRI data (Fogelson et al., 2011). It was found that the accuracy of classification using ST-embedded fMRI data (i.e. entire-trial ST embedding) is higher than using individual, temporally distinct spatial-only observations (In the current study the aforementioned technique is referred to as single TR observations). ST embedding was also investigated for another type of stimulus classification in (Rao, Garg, & Cecchi, 2011) applying the same methods discussed by Fogelson and colleagues (Fogelson et al., 2011). Another study investigated the variability of temporal dynamic classification performance across single TR observations within the slow event-related trails (Kohler et al., 2013). Their timepoint-by-timepoint MVPC showed that the peak of classification accuracy was around the peak of region-average HRF; in some regions prior to and in some regions after the region-average HRF peak. However, it was not clear whether all of the temporal dynamics of a voxel activity within a trial carry stimulus specific information.

On the other hand, most of the multivariate pattern recognition based studies applied to fMRI data, modeling the pairwise relationship between the brain activity at separate time points and the experimental condition. Although considering the shape of HRF for modeling brain activity, they assign the same stimulus label to the dynamic brain response that is changing over time. The temporal dynamics of the BOLD signal to stimuli of different classes, and within different brain regions are very likely different (Chu et al., 2011; Kohler et al., 2013). Not considering such these differences may reduce the sensitivity of the classifier and reduce the decoding accuracy.

ST feature selection of fMRI finds the time points that carry the highest condition specific information for MVPC (Choupan et al., 2014). The effect of ST feature selection initially was tested on a block-design experiment (Choupan et al., 2014). We (Choupan et al., 2014) found that ST feature selection could improve prediction accuracy even on a block-design experiment. To study ST feature selection thoroughly, however, a slow event-related design is preferred because it provides a “cleaner” temporal pattern in which the neural response is less affected by the temporal overlapping of consecutive stimuli responses that occurs in block or rapid event-related design. Particularly when the BOLD signal is allowed to return to baseline before the next stimulus is presented. In comparison to block designs, by randomizing condition/stimuli order, slow event-related experiments minimize effects of strategy expectation and cognitive set (which affect the temporal dynamics) (Pilgrim, Fadili, Fletcher, & Tyler, 2002; Strayer & Kramer, 1994). In addition, slow event-related designs reduce the neuronal habituation that has been shown to alter the results of MVPC in block design experiments (Choupan et al., 2014; Mourao-Miranda, Friston, & Brammer, 2007; Sapountzis, Schluppeck, Bowtell, & Peirce, 2010).

Following the previous works, this study is based on the hypothesis that by embedding the temporal dynamics provided by fMRI into the process of multivariate brain pattern recognition, more information contained in the BOLD signal can be utilized compared with single TR methods, leading to a potential improvement in the prediction performance. In particular, we predicted that not all of the temporal dynamics of voxels within the trial are informative for the MVPC, and that a shorter *sequence* of time points might still possess the most discriminative activity across stimulus conditions, possibly around the peak of the HRF (Akama, Brian Murphy, Shimizu, & Poesio, 2012; Formisano, De Martino, & Valente, 2008; Kohler et al., 2013). Therefore, this study presents an investigation of the MVPC performance of ST feature selection using different sequences of ST combinations.

On a dataset acquired with a CMRR multi-band EPI pulse sequences at a high temporal resolution of 0.568 seconds in a binary slow event-related design (inter-stimulus interval of ∼25 seconds), using stimuli from the classic Haxby et al. (2001) experiment, we assessed the prediction accuracy of 990 ST combinations. Our findings show that on average, ST feature selection led to improved classification performance. Furthermore, the discriminative power increased when time points around the peak of the HRF were included in the ST combination.

## 2. Materials and Methods

### 2.1. Participants

Four right-handed healthy adult volunteers (ages 28, 30, 31, and 32; two of them were females) participated in this study. None had a medical history of psychiatric disorder, as assessed by self-report. Written informed consent, approved by the University of California, Los Angeles Institutional Review Board, was obtained from each participant prior to the experiment. The heart rate and skin conductivity of participants were recorded and monitored throughout the experiment.

### 2.2. Experimental design

The task paradigm was implemented in MATLAB (Mathworks, Inc.) using the Psychophysics Toolbox, Version 3.0 (Brainard, 1997). Stimuli were projected onto a screen behind the scanner bore which participants watched through a mirror installed on the head coil. Participants engaged in six 11-minute fMRI scans. Each run consisted of twenty slow event-related trials, yielding a total of 120 trials per participant. During each fMRI run, participants viewed 10 pictures of human faces (five were females) and 10 pictures of houses. The pictures, which were borrowed from publicly available stimulus set used in the Haxby and colleagues paper (Haxby et al., 2001), were displayed in random order, different for each subject and trial. In each trial, participants viewed a single stimulus picture for 500 ms, which was always followed by an inter-stimulus interval of 25s (the stimulus onset times were jittered at each trial to avoid anticipatory brain activations). In each run, three random trials were followed by their content photographed from different angle. We asked the participants to perform a one-back repetition detection task. The participants were provided with an MRI compatible button box to indicate their responses. In the case of similar consecutive trials (identical re-oriented pictures), participants were instructed to press the right button of the button box, and the left button for the dissimilar pictures. The similar trials, which were employed solely to ensure that subjects remained awake and engaged, were excluded from the analysis. We chose a long Inter Stimulus Interval (ISI) of 25 seconds in considering the standard double gamma HRF function characteristics (Friston et al., 1994) that requires ∼25 seconds for BOLD signal to get back to the baseline after observing a stimuli (Cohen M. S., 1997).

### 2.3. Data acquisition

We acquired images were using the Siemens 3T Tim Trio scanner with a 32-channel head coil at the Staglin Center for Cognitive Neuroscience at the University of California, Los Angeles. The functional images were acquired with a CMRR multi-band EPI pulse sequences C2P, 010b (Auerbach, Xu, Yacoub, Moeller, & Uğurbil, 2013; Moeller et al., 2010; Setsompop et al., 2012; Sotiropoulos et al., 2013; Xu et al., 2013) with multiband acceleration factor of four, and phase encoding direction acceleration factor of 3 (referred to as integrated-Parallel Acquisition-Techniques or iPAT factor in Siemens terminology) yielding a net acceleration of 12 (4×3). In addition, in-plane rotation was set to 180-degree, TR = 0.568 s, TE = 0.3 s, flip angle = 40°, 40 slices, 3×3×3mm, FOV = 192×192 mm (axial acquisition) covered the whole brain. SBRef data was collected, but not utilized for pre-processing. No field map data was collected. This setting resulted in 45 TRs per trial. A high-resolution structural T1-weighted MPRAGE was acquired for each participant (176 sagittal slices, 0.97×0.97 mm in-plane voxel resolution, 1 mm slice thickness, matrix size = 256×256, FOV = 250×250×176 mm, TR = 1.9 s, TE = 2.26 s, flip angle = 9°).

### 2.4. Data pre-processing

We pre-processed the functional images using Statistical Parametric Mapping software (SPM8; http://www.fil.ion.ucl.ac.uk/spm). Because we used multi-band acquisition, no slice-timing correction was applied (Glasser et al., 2013). Each fMRI volume was first realigned to its mean image using the 4^th^ degree B-spline interpolation for head motion correction. The anatomical volume was segmented to gray matter, white matter, and cerebrospinal fluid. We registered the functional data from each run to the anatomical volume, then spatially normalized the data into standard stereotaxic space with voxel size of 2×2×2 mm^3^, using the Montreal Neurological Institute (MNI) template. Warping to MNI was performed to assure that the input data for each subject has the same size/dimension across subjects.

As recommended by Kohler (Kohler et al., 2013; Misaki, Luh, & Bandettini, 2013), we applied no spatial smoothing.

At each separate fMRI run, we linearly-detrended the voxels time course to reduce the effects of signal drifts during the course of fMRI experiment, then, we normalized the detrended voxels time course across the entire run to zero mean and unit variance across observations.

Field map data was not collected at the time of data collection, and field map inhomogeneity distortion correction was not performed. Regardless, visual inspection showed that the fMRI images were registered to structural image with minimal distortion.

### 2.5. Region of interest

Our selection of the regions-of-interest (ROIs) was based on prior knowledge. Previous studies have shown that inferior temporal (IT) gyrus exhibits category-specific responses during perception of faces (in FFA) or scenes (in PPA) (Kriegeskorte et al., 2008; Ranganath, DeGutis, & D’Esposito, 2004). Therefore, our analysis was restricted to bilateral IT defined as in the AAL atlas in the WFU_PickAtlas MATLAB software toolbox (http://fmri.wfubmc.edu/software/PickAtlas). Functional images, and the ROI mask, were defined in the MNI space, and the derived mask was applied to the preprocessed functional images. In total there were 7547 voxels in IT, using WFU_PickAtlas.

Training the classification algorithms on total number of voxels in IT in spatiotemporal form was computationally intensive. Therefore, we decided to perform spatial feature selection. Random Forest (RF) (Breiman, 2001) was utilized as a spatial feature selection method to further reduce the size of the already masked data, discarding the voxels that do not improve category specific classification. RF feature selection calculated the voxels importance for the training data. Voxels importance was calculated based on the mean error of bootstrap tree samples in the forest. During the bootstrapping procedure, the voxel is randomly permuted in the Out Of Bag (OOB) cases. The aim of this permutation is to eliminate the existing association between voxels and the stimuli, and then to test the effect of this elimination on the RF model among trees built on these bootstrap samples. A voxel is considered to be in a strong association with the stimuli if the mean error decreases.

For each subject, the spatial feature selection was applied on IT voxels (containing 7547 features), on 100 trials as training samples. 1000 trees were utilized to train the RF model. After training, voxels in the top 1% of maximum OOB importance were selected, resulting 115 voxels for each subject. All subjects were registered to MNI space, which results in same number of voxels after feature selection.

### 2.6. Spatiotemporal data representation

For each trial of fMRI, all possible ST combinations were defined from the data acquired at all *N =* 45 TRs. In another word, for investigating the most informative temporal features, the entire hemodynamic response temporal domain was searched for voxels/features picked by RF feature selection. Assuming that the combination should have at least 2 time points of voxels activity, and the combinations should be continues and in ascending orders according to time, this representation led to obtaining in total 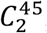, 990 ST combinations for each fMRI run. The ST observations for each trial were defined as

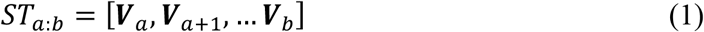

where ***V***_*i*_ were the BOLD signals at volume *i* ∈ {1,2, …,45}, *a* = 1, 2, …, *N* − 1, *b* = 2, 3, …, *N and a* < *b*. Therefore, each *ST_a:b_* was the result of concatenating voxels activity in ***V**_a_*… ***V**_b_*. Utilizing such a concatenation routine, the temporal information was embedded together with the spatial information, forming an ST observation. The prediction accuracies of entire cases were explored to investigate the informative duration of BOLD signal for classification relative to the stimulus onset. The concatenation process in ST is illustrated in Figure 1.

**Figure 1.**
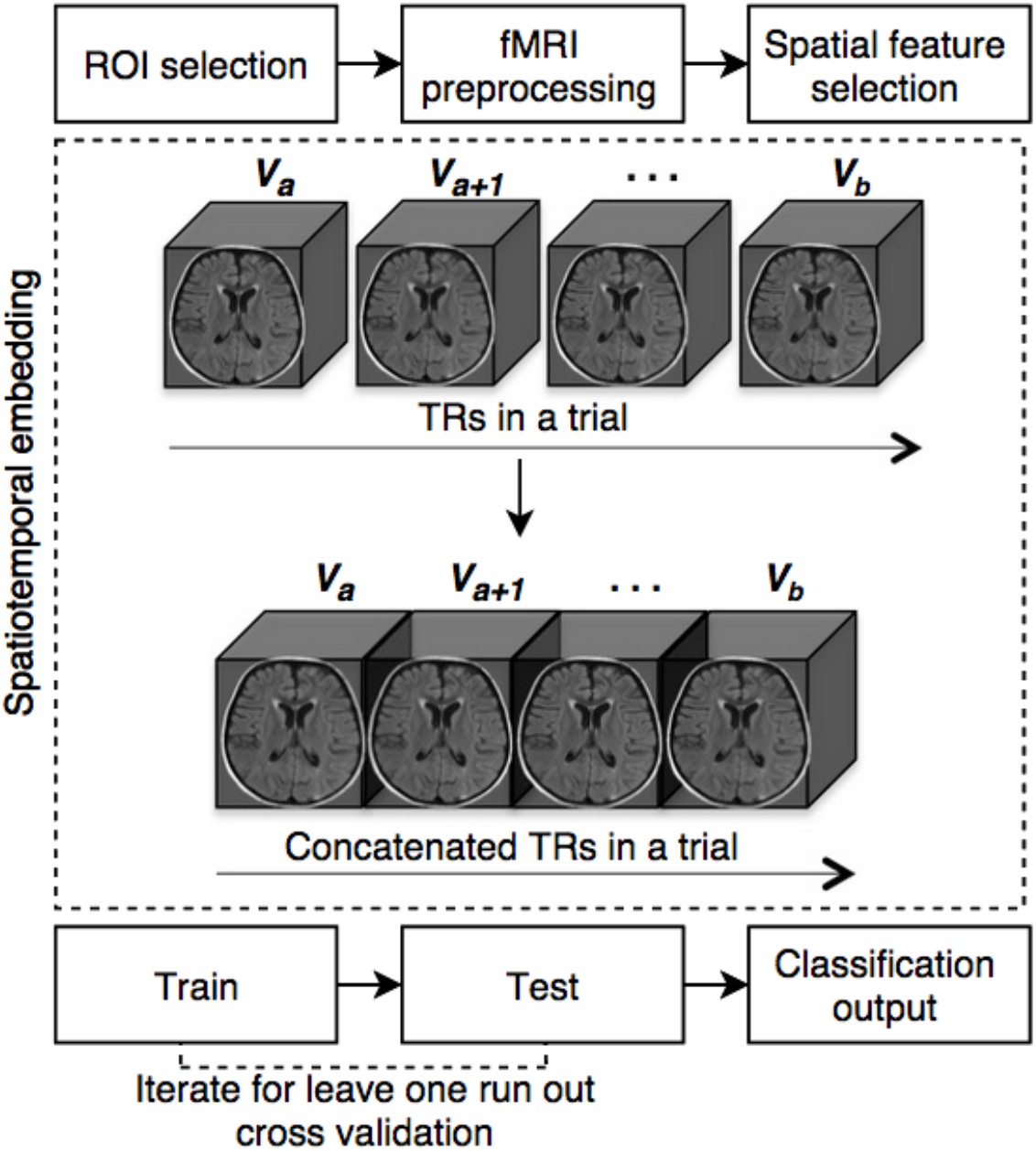
Graphical illustration of data processing. The top row indicates the preprocessing steps employed in this study. Spatiotemporal embedding is shown in the dotted box, which involves concatenating the fMRI volumes, a through to b. Each 3D cube is a symbolic fMRI volume. The bottom row illustrates the learning steps.

In a previous slow event-related decoding study, which investigated the temporal domain effect on MVPC (Kohler et al., 2013), the inter-stimulus interval was around 11 seconds. Therefore, an extra examination was performed to validate the effect of ST embedding on overall prediction accuracy of all subjects in this study, using only the first 11 seconds after the stimulus onset.

### 2.7. Single TR data representation

For comparison purposes, we used single TR spatial observation (utilizing the spatial information acquired during one Time to Repeat or (TR)). The maximum accuracy of single TR observation in slow event-related fMRI was reported to be around the peak of HRF at ∼5 seconds, but with a small jitter across regions (Kohler et al., 2013). The HRF peak has also been found to be jittered across people (∼ 4 to 7 seconds) (Handwerker, Ollinger, & D’Esposito, 2004). Therefore, single TR classifications were performed for all TRs in the above range (1 second before and 2 seconds after the HRF peak) and only the highest performances were reported. This approach assured that our ST combinations were compared with the highest performance of the single TR approach. It should be noted that the high temporal resolution of the acquired fMRI data allowed us to perform this rigorous investigation.

### 2.8. Pattern classification

Multivariate brain pattern recognition was performed using the Princeton MVPA toolbox (http://code.google.com/p/princeton-mvpatoolbox/). Support Vector Machine (SVM) is a widely used classifier in the field of neuroimaging and MVPC applications (Bode & Haynes, 2009; Kohler et al., 2013; Mourao-Miranda et al., 2007; Rao et al., 2011; Ritter, Hebart, Wolbers, & Bingel, 2014; Waskom, Kumaran, Gordon, Rissman, & Wagner, 2014). Douglas et al. found that RF outperforms SVM in a binary classification of belief vs. disbelief using fMRI data (Douglas, Harris, Yuille, & Cohen, 2011). Hence, in this study the two classification algorithms RF and SVM were employed and compared. In addition, the two classifiers allow the extraction of feature weight vectors, indicating discrimination power. All analyses were performed using MATLAB software (V. 8.5 Mathworks, Inc.). MATLAB-based tools for RF (Jaiantilal, 2009) and linear SVM (Fan, Chang, Hsieh, Wang, & Lin, 2008) were utilized. For SVM analyses the regularization parameter that controls the trade-off between model fitting error and classification accuracy, was set to 1 (Waskom et al., 2014). A leave one run out cross validation (LOROCV) scheme was employed (Pereira, Mitchell, & Botvinick, 2009). In experiments on both single TR and ST combinations, the fMRI data were divided into training and test sets, training the classifier using five runs and testing on the sixth run. This test was repeated 6 times, with each of the different runs serving once as a test set. Finally the prediction accuracies were reported to quantify how accurately the classifiers were able to distinguish between faces and houses.

SVM (Burges, 1998; Vapnik, 2000) was employed in similar temporal investigation decoding studies (Kohler et al., 2013; Mourao-Miranda et al., 2007). The learning process of SVM classifier finds the maximum-margin hyperplane that separates the training data observations according to the class they belong (faces or houses in this study). This hyperplane is orthogonal to the direction along which the training observations of both classes differ most.

Linear SVM training outputs a set of weights, one for each feature, whom their linear combination predicts the value of stimuli categories. This weight vector allows investigation on the discriminating power of features across stimulus categories. The directions of the weight vectors are perpendicular to the separating hyperplane. A feature with a positive weight value means that the feature has higher activity (discrimination power) for stimuli 1 than stimuli 2 in the training examples. The weight patterns were reconstructed according to (Haufe et al., 2014) by applying the following algorithm:

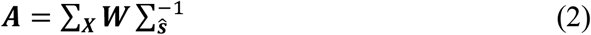

where, ***A*** is the reconstructed pattern, ***W*** is the weight vector, ∑_***X***_ is the n-by-p covariance matrix of the data (with n voxels and p samples), and 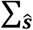 is the source covariance, defined as ***W**^T^* × ***X***.

RF is an ensemble classifier that employs *decision trees* as base learners (Breiman, 2001). In this algorithm, training set observations is resampled (random redistribution, with replacement) multiple times using bootstrap technique to produce multiple training subsets. Decision trees are then created from each training subset, until all ensembles of trees have been created. For predicting the label of an unseen testing observation at each tree, the data is feed to the root of the tree, and goes down the tree following the splits and falls into a terminal node. Each tree outputs the label in the terminal node. Final predictions are assigned based on the majority voting on trees label decision. 1000 trees were utilized for training the RF model, and the number of trees was selected based on the stability of the OOB error rate to an asymptotic plateau.

### 2.9. Performance evaluation

Two criteria were employed to evaluate the performance of the classification at each cross validation run: the overall prediction accuracy, and sensitivity to each stimulus, or recall. Overall accuracy was calculated as the percentage of correctly classified trials at each testing step. Sensitivity or true positive rate was measured as

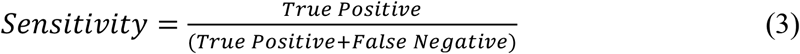

that is the number of correctly predicted positive instances over all positive instances. When measuring sensitivities, the category of interest was considered as positive instance. When comparing different techniques, the mean of derived cross-validation runs was reported. We obtained the same results when looking at specificity (or precision). Therefore, we only reported sensitivity for readability.

### 2.10. Analysis of single TR combinations results

In total, the ST and single TR methods were utilized with SVM and RF classifiers to perform the analyses: ST-SVM, ST-RF, Single TR-SVM, and Single TR-RF. The best performance of the entire ST combinations was compared with the best single TR approach around the peak of HRF to investigate if ST embedding can improve the classification. Then, the prediction accuracy of all ST combinations were plotted and mapped to explore the most discriminating temporal duration for decoding. For all above cases RF and SVM were compared with each other. The temporal duration of top performed ST combinations were plotted to investigate which classifier benefits more from temporal embedding, the longer the ST combination of top performed classification is, indicates that the classifier benefits more from the temporal information compared with the other classifier.

All aforementioned analyses were performed on participants 1. As a result of deriving the important temporal duration across all ST combinations in participant 1, the duration was employed to analyze best performance across all participants to investigate if a shorter inter-stimulus interval, which is similar to previous work (Kohler et al., 2013), could still provide higher prediction accuracy compare with Single TR technique.

### 2.11. TR influence index

This study investigated the influence of data acquired at each TR over the course of fMRI, relative to the stimulus onset, on the classification performance for each stimulus category across all 990 ST combinations. Firstly, the most discriminating ST features (*i.e.*, the BOLD signal acquired at a TR in a voxel) in each ST combination were determined from the training results. When SVM was employed, on each stimulus side of the hyperplane, the selected features were the top 1% ST features with the largest reconstructed weight value. When RF was employed, the top 1% ST features with the largest OOB importance were selected. Secondly, the presence of a given TR in all selected ST features was counted as an indication of its influence on the classification performance. Thirdly, the presence of a given TR was normalized by the total number of times (*P*) that TR was presented in all 990 ST combinations. *P* is calculated as *P* = *T*(*N* − *T* + 1) − 1, where *T* is the serial number of the given TR, ranging from 1 to 45, and *N* is 45, the total number of TRs. This study calls this normalized value *TR influence index*.

BOLD signals measured via fMRI are very slow. A TR with high influence in a ST combination window affects its neighbors not to be selected, until that TR is out of the ST window. Therefore, for better representation the results were overlaid with the maximum TR influence for tri-seconds interval. This time interval almost mimics the temporal resolution of conventional fMRI sequence.

## 3. Results

### 3.1. Brain decoding based on spatiotemporal features versus spatial-only single time point technique

The ST embedding based techniques resulted in higher cross-validated prediction accuracy in comparison to single TR techniques. This improvement was consistent across separate stimuli categories for the two studied classification methods. Table 1 showed that on average, across six runs RF (over cross-validate accuracy of ST and single TR techniques were 81.66 and 69.16, respectively) outperformed SVM (over cross-validate accuracy of ST and single TR techniques were 78.33 and 58.33, respectively) overall, and in separate stimuli specific evaluation. When looking at the stimulus-specific results (Figure 2B-C), ST techniques showed higher sensitivity (19-25% higher). In particular, ST-RF sensitivity to independent stimuli were always 90% or higher. Single TR SVM, which is among the most popular techniques (Kohler et al., 2013; Mourao-Miranda et al., 2007), performed slightly higher than chance. However, by using ST-SVM, sensitivity to independent stimuli were 80% or higher.

**Figure 2.**
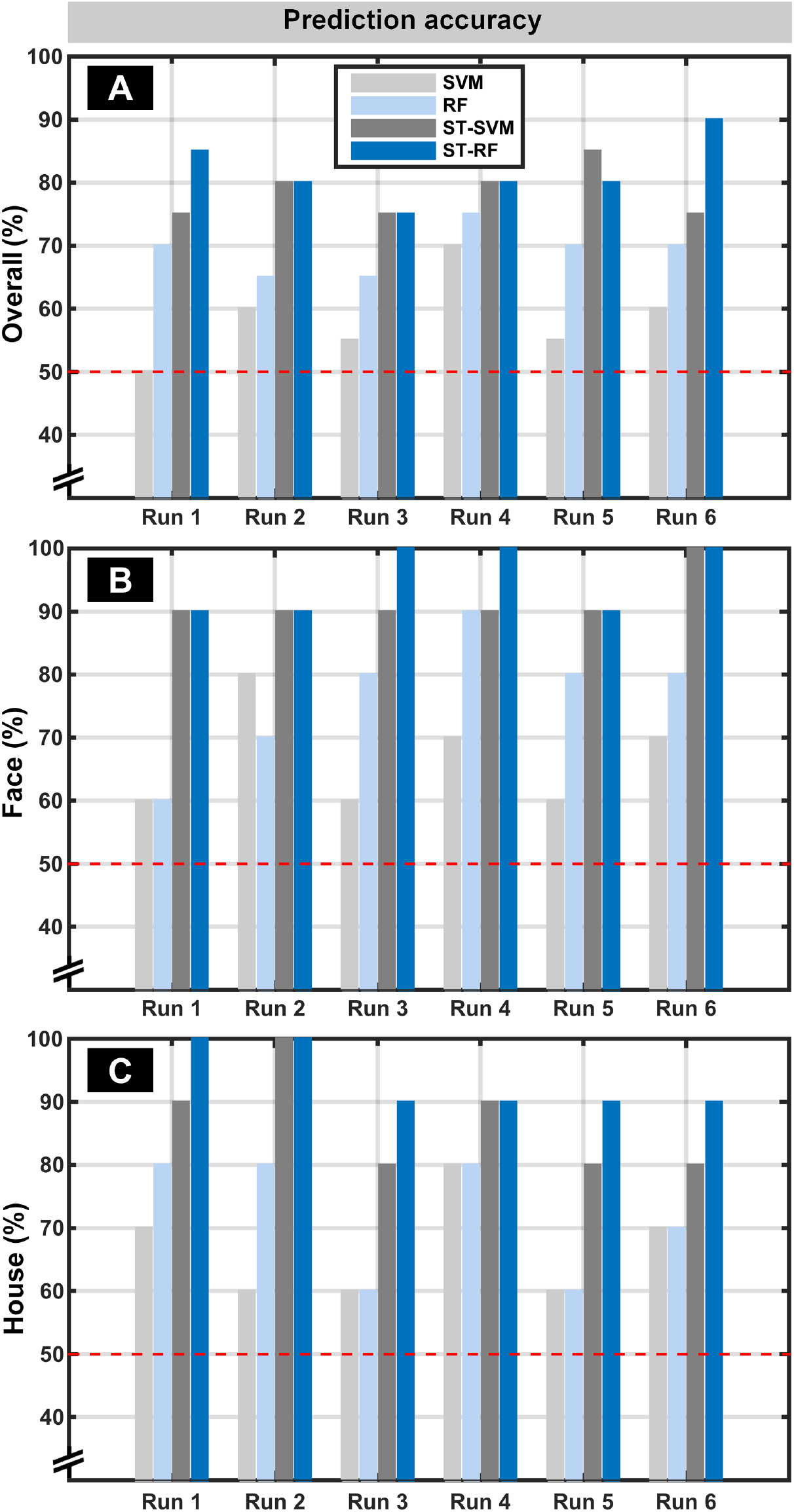
Best performance of each technique across six runs. **(A)** Overall accuracy of single TR Support Vector machine (SVM) and Random Forest (RF), together with SpatioTemporal SVM (ST-SVM) and ST-RF. For single TR, the canonical Hemodynamic Response Function (HRF) peak with 1 second before and 2 seconds after the peak was considered and the highest performance was reported. For ST techniques, the highest achieved prediction accuracy is illustrated. The red line indicates the chance level accuracy (50%) for faces versus houses classification. Sensitivity in detecting faces and houses are illustrated in **(B)** and **(C)**, respectively.

**Table 1.** Average performance of each technique across six runs. Average performance of single TR Support Vector machine (SVM) and Random Forest (RF), together with Spatio-Temporal SVM (ST-SVM) and ST-RF (rows) across six cross-validation run. Columns from left to right present the overall performance, accuracy on predicting Face stimuli, and House stimuli, respectively.

|  | Overall | Face | House |
| --- | --- | --- | --- |
| SVM | 58.33 | 66.66 | 66.66 |
| RF | 69.16 | 76.66 | 71.66 |
| ST-SVM | 78.33 | 91.66 | 86.66 |
| ST-RF | 81.66 | 95 | 93.33 |

When looking at the best prediction outcome of different classifiers, we noticed that RF outperformed SVM in most of the instances. The superiority was consistent across all runs and categories, where RF results were almost always higher than SVM and ST-RF results were always equal or higher than ST-SVM. It should be noted that here only the highest achieved prediction accuracies and sensitivities were reported. Therefore, the reported results in Figure 2 A, B and C are not from the same ST combination. It can be seen that the overall accuracy is mainly lower than the highest accuracy achieved in detecting either faces or houses.

Prediction accuracies of all possible ST combinations are demonstrated in Figure 3. In this figure, all the 990 ST combinations are ordered next to each other in a way that the early results are the ST combinations where the early time points are included in the ST time window, and each immediate neighboring result is from the ST window expanded to the next time point until it reaches to time point = 45. Note that by using ST embedding technique, not all ST-RF combinations outperformed single TR-RF, and the classification performance in many of the combinations are even lower than chance. While ST embedding improves prediction accuracy in ST-RF compared to the Single TR-RF (around canonical HRF peak), the improvement highly depends on the choice of ST combination. For example, in ST-RF the accuracy was higher than Single TR-RF mainly when early TRs in the first third of the time interval after the stimulus onset were included. However, in the later combinations, which associated with combinations containing the last third TRs in the trials, the performance of ST-RF was lower than Single TR-RF. No consistent pattern was observed in the results of ST-SVM.

**Figure 3.**
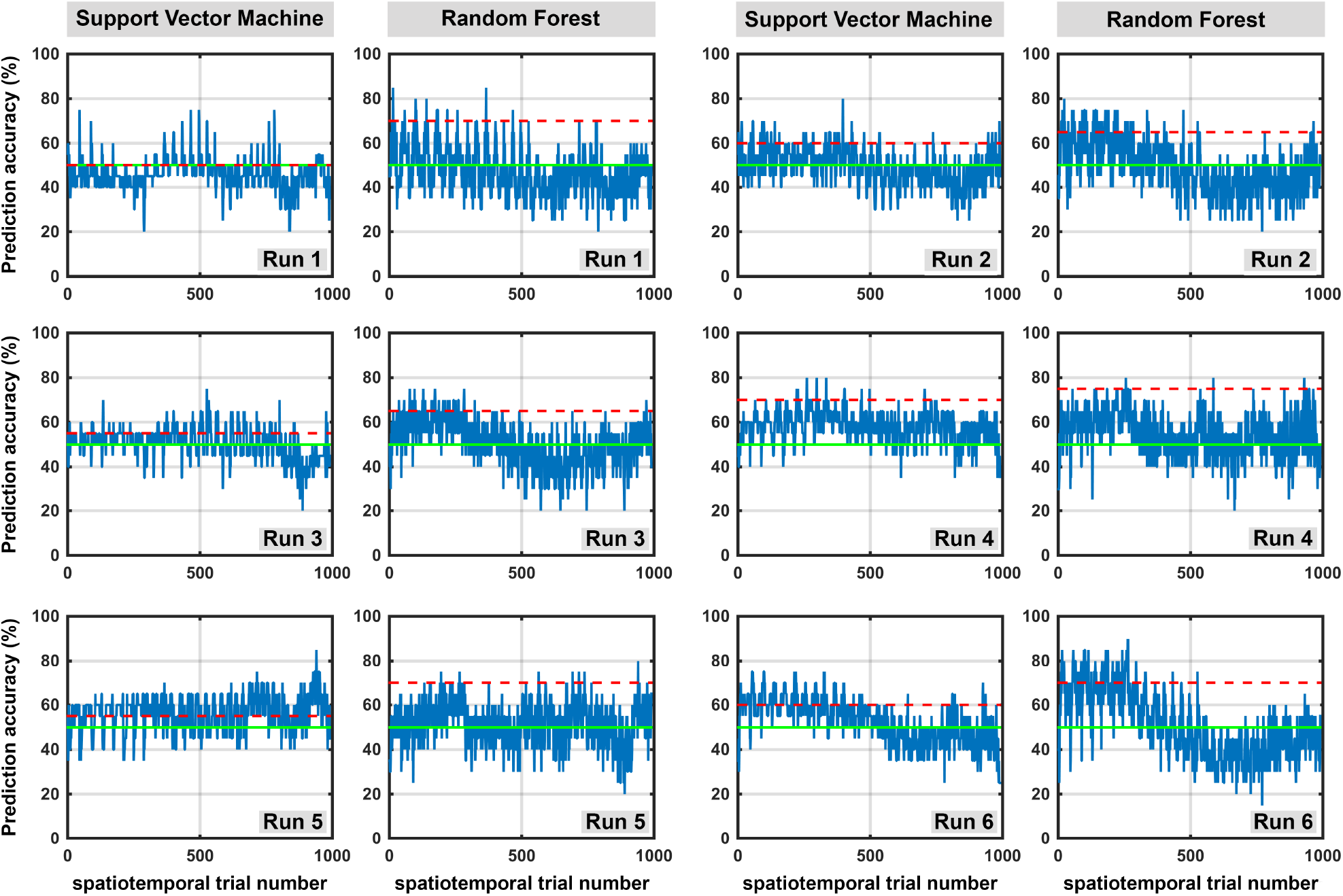
Overall prediction accuracy of all possible spatiotemporal combinations across six runs. Each box shows the prediction accuracy of 990 possible SpatioTemporal (ST) combinations out of 45 TRs. The odd and even columns represent prediction accuracies using Support Vector machine (SVM) and Random Forest (RF) classifiers, respectively. Green line indicates the prediction chance (50%), and red line represents performance of the single TR technique (around canonical HRF peak) of that run. The x-axis shows the 990 ST combinations beginning with 1^st^ TR (i.e. TRs: 1-2, 1-3, 1-4, …, 1-45) followed by all combinations starting with 2^nd^ TR (i.e. TRs: 2-3, 2-4, 2-5, …, 2-45) and so forth. The 990^th^ ST trial includes the 44^th^ and 45^th^ TRs.

Figures 2 and 3 showed that across all runs Single TR-RF performs better than Single TR-SVM. Single TR with RF classifiers was even as high as ST-SVM in most cases, but not better than the best performing ST-SVMs.

### 3.2. Investigating discriminative temporal duration

In order to explore the most informative temporal duration using ST, all ST trials were mapped in Figures 4 and 5. The two figures reflect a heatmap of the result space, and show where, in temporal duration, high informative spatiotemporal combinations are centered. The color distribution represents the strong and weak prediction accuracies. Using ST-RF, a trend in prediction accuracy was observed (Figure 4). The high accuracy was mainly concentrated in the left side of the maps, which is associated with ST combinations that started at the early time points from stimuli onset. A noticeable drop in prediction accuracies was seen when the beginning of ST-RF was 6s or later. In some runs or categories, most of 45 TRs were included in the ST leading to high accuracy, but their accuracy never exceeded the ST combinations *ST*_5:20_, including TRs from ∼2 seconds to ∼11 seconds.

**Figure 4.**
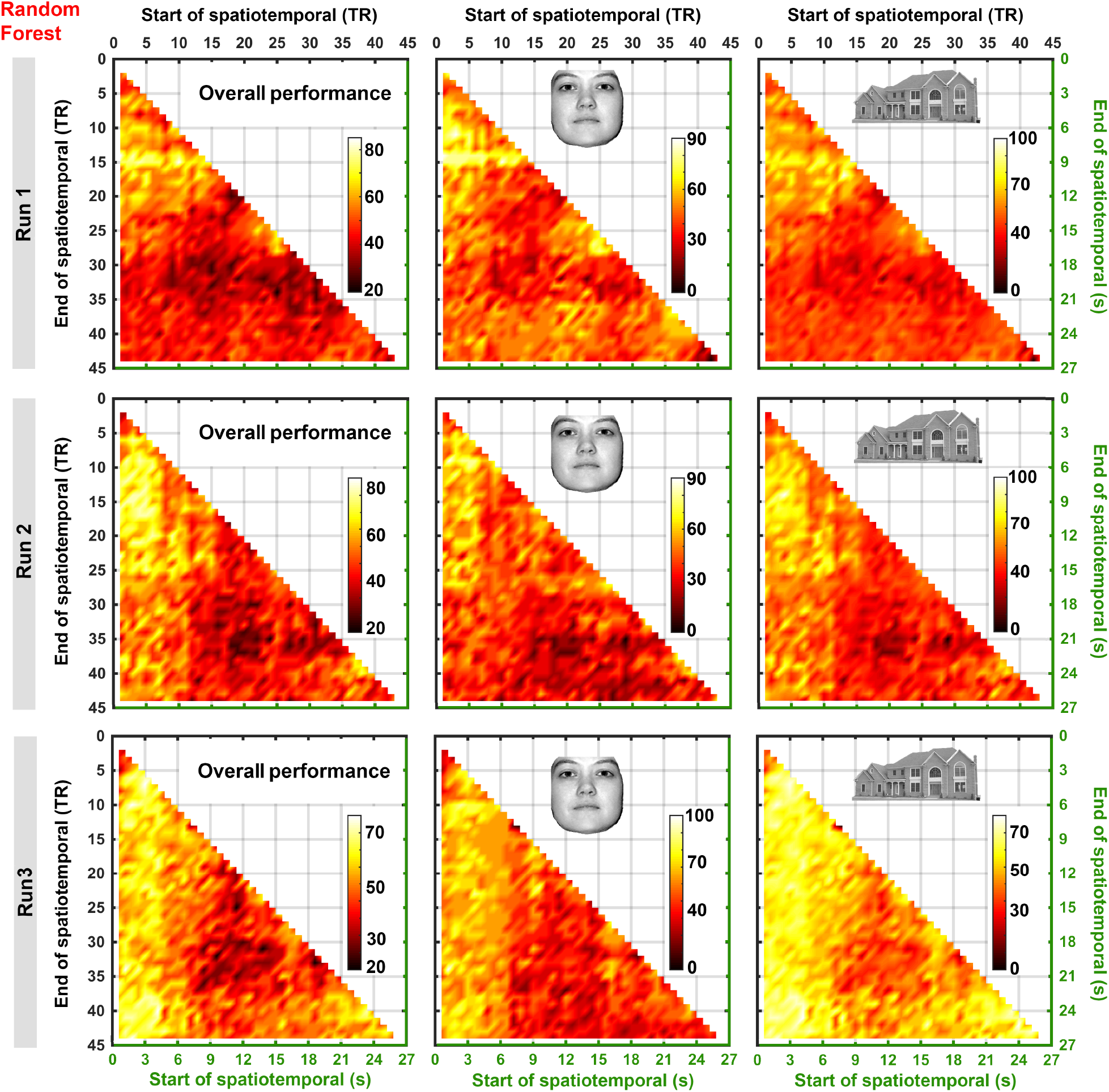
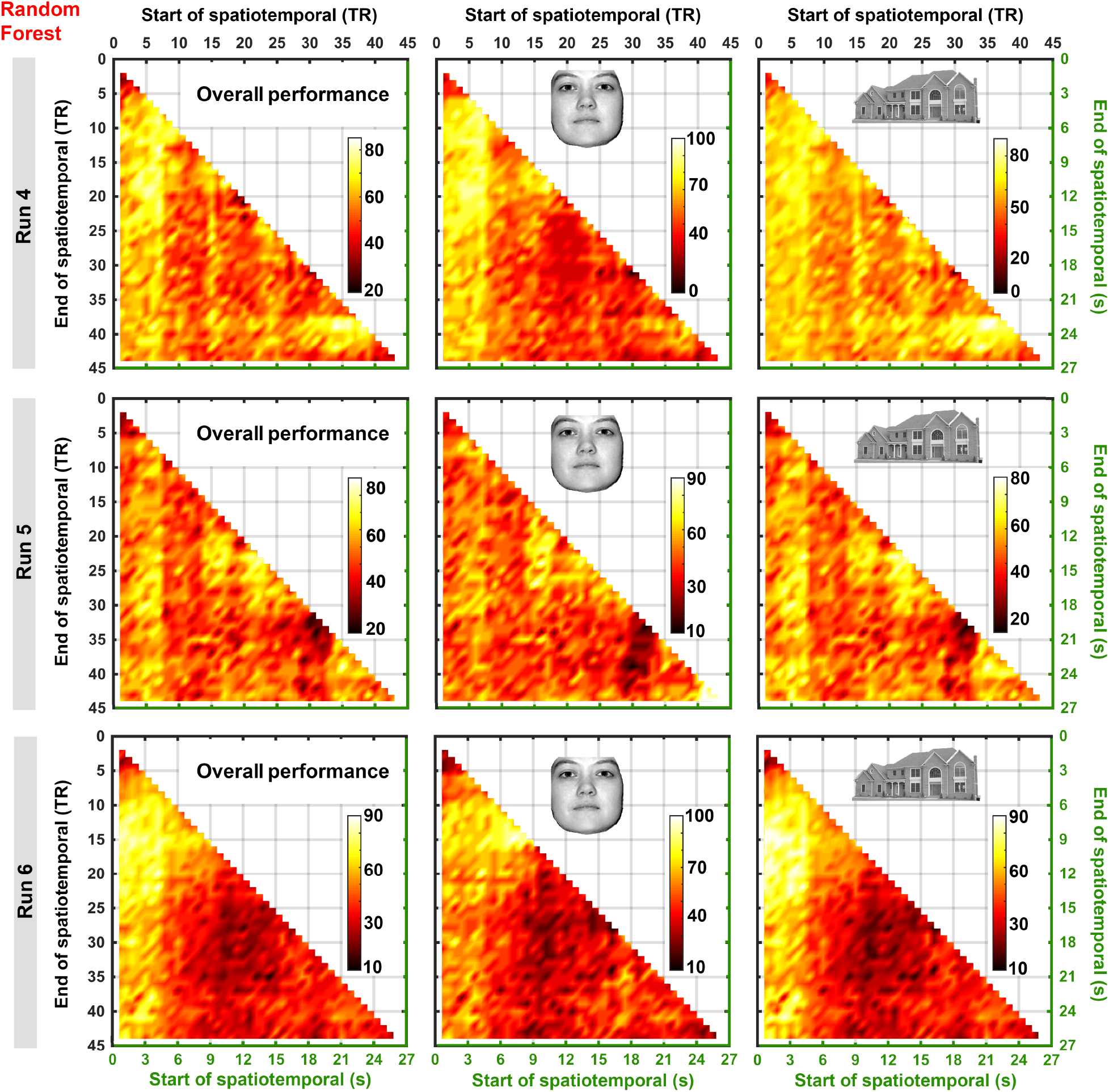
Prediction accuracy of all possible spatiotemporal combinations across six runs using Random Forest classifier. First column of each hot map shows the overall prediction accuracy of 990 possible SpatioTemporal (ST). Second and third columns of each hot map show the sensitivity in predicting face and houses, respectively. X- and Y-axes indicate the start and end time of the ST, respectively. The precise time (in seconds) of each ST combination is shown in green axes. For example, point [2,18] in these maps is the ST combination that starts from TR of 2 (on X axis) and ends at 18 (on Y axis). The columns represent overall prediction accuracies and sensitivity to faces and houses, respectively. Different color map ranges were used for each map, to assist visual inspection of most informative intervals.

**Figure 5.**
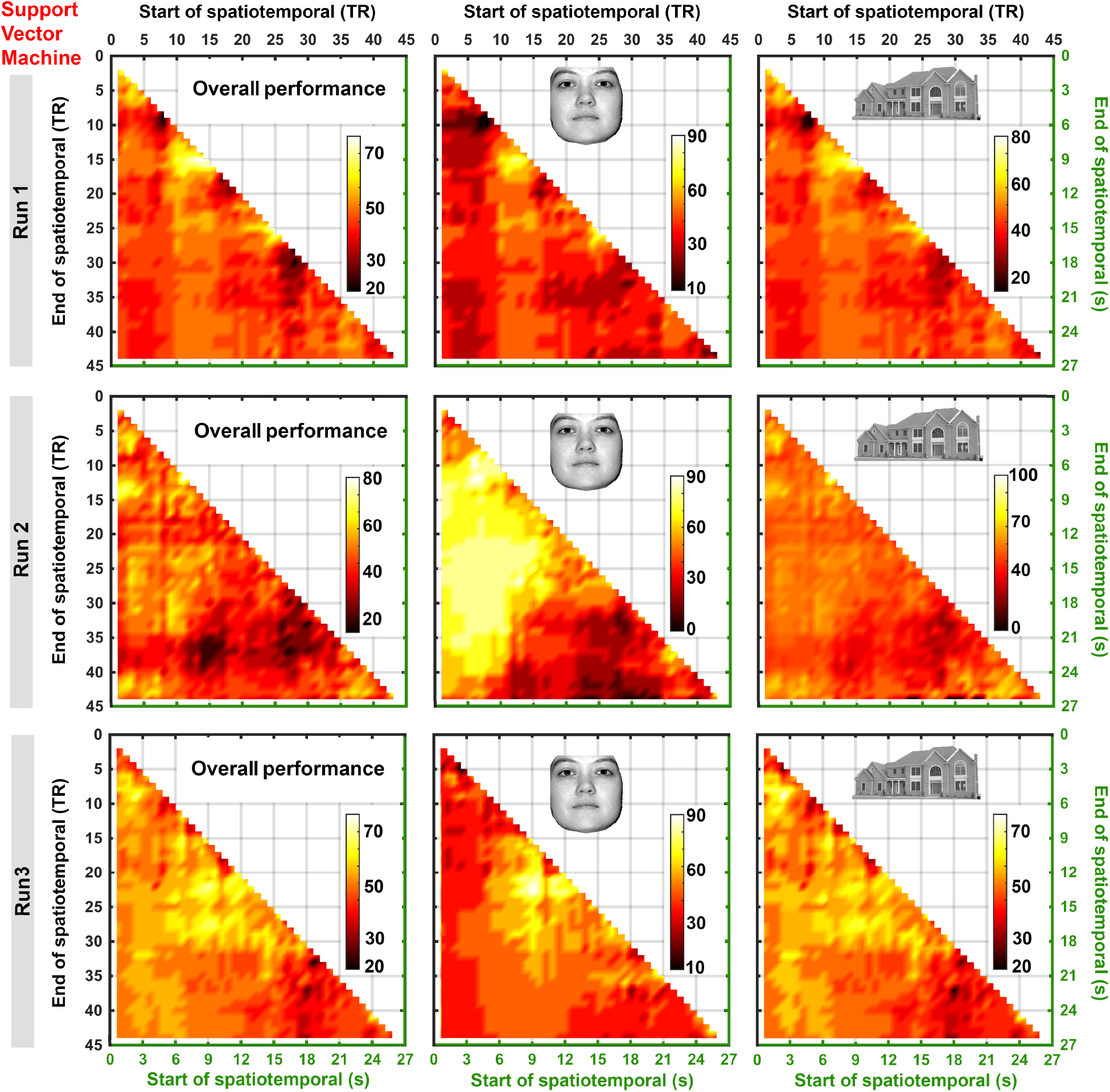
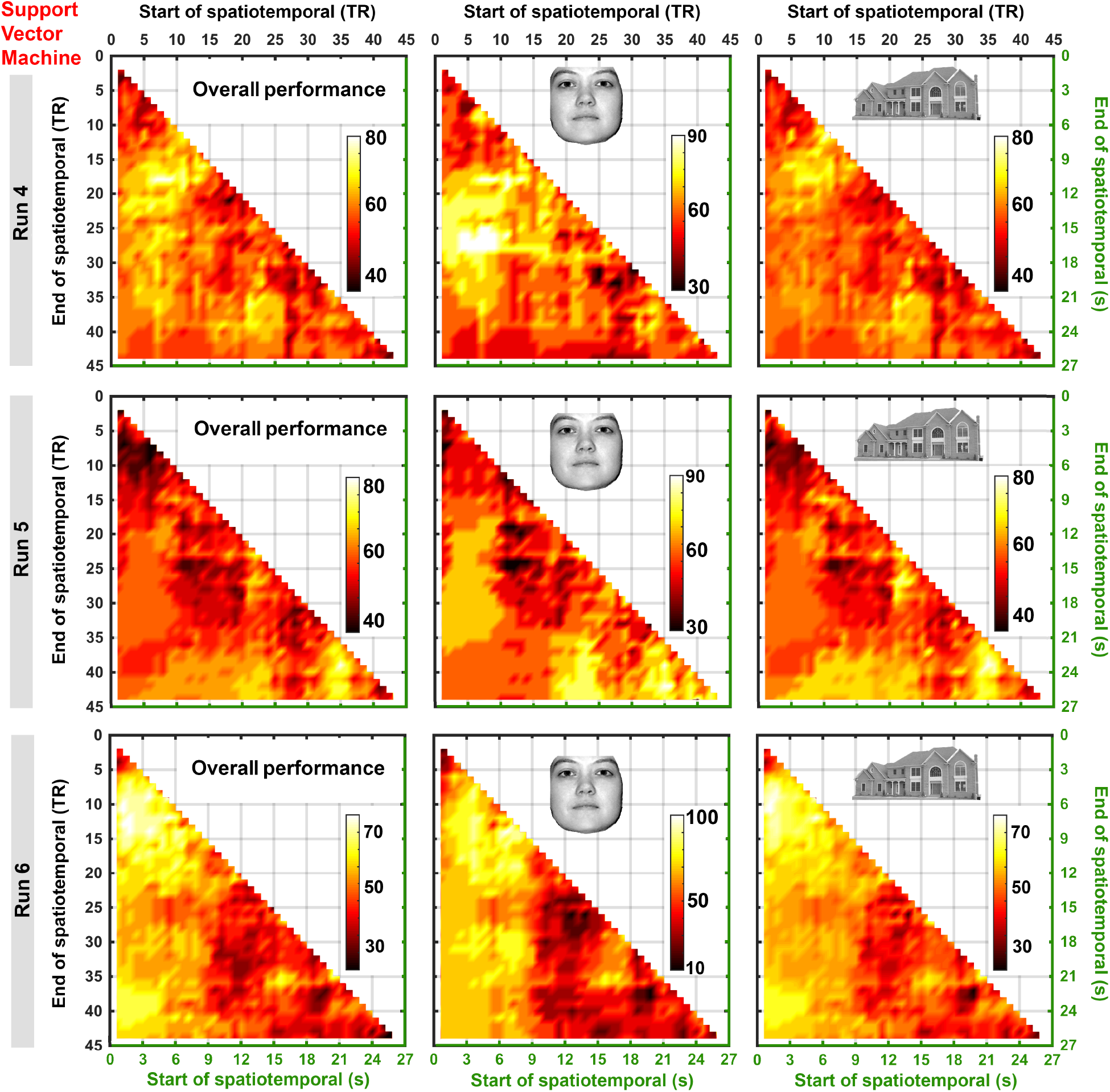
Prediction accuracy of all possible spatiotemporal combinations across six runs using Support Vector Machine classifier. First column of each hot map shows the overall prediction accuracy of 990 possible SpatioTemporal (ST). Second and third columns of each hot map show the sensitivity in predicting face and houses, respectively. X- and Y-axes indicate the starting and ending time of the ST, respectively. The precise time (in seconds) of each ST combination is shown in green axes. For example, point [2,18] in these maps is the ST combination that starts from TR of 2 (on X axis) and ends at 18 (on Y axis). The columns represent overall prediction accuracies, sensitivity to faces and houses, respectively. Different color map ranges were used for each map, to assist visual inspection of most informative intervals.

Across all runs and categories, the prediction accuracy of ST combinations with end points smaller than TR 5, e.g. *ST*_1:4_ or *ST*_2:3_, were around chance. These ST combinations can be seen in the top left corner of the maps in Figure 4. As soon as the ST combination started to include the preceding TRs, increased prediction accuracy was observed. In quite a few instances, an increase in prediction accuracy was observed in the ST combinations including late TRs, which are around 20 seconds after stimulus onset. This increase can be visualized as the hyperintensity patch in the right bottom corner of the maps. The importance of individual TRs was further investigated, as illustrated in Figure 6 and 7.

**Figure 6.**
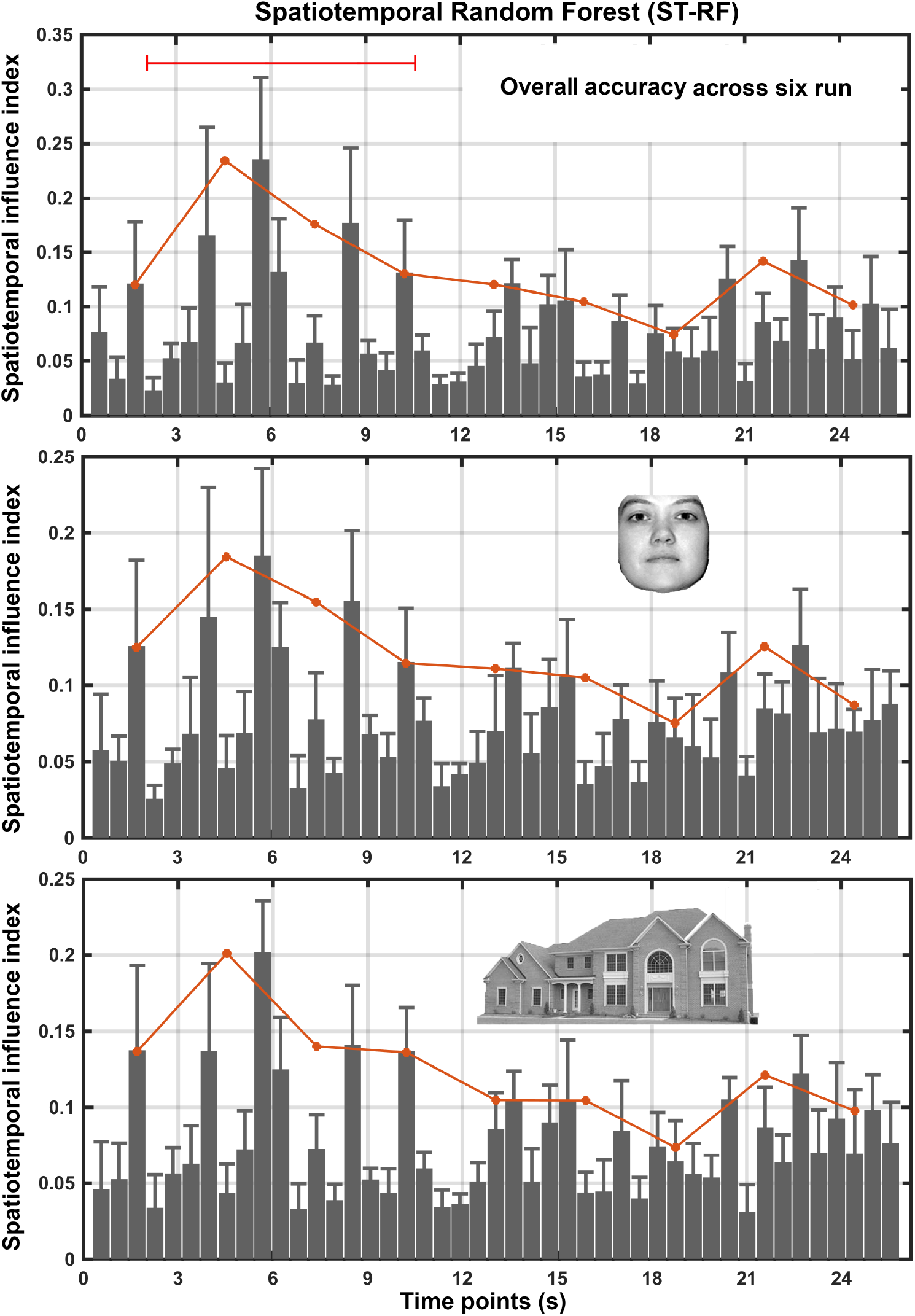
Influence of each time point using Random Forest (RF). Bars indicate the mean and standard deviation of TR influence index across six runs. TR influence index represents the number of times where each time point was chosen as the most informative time point of the ST combination, normalized over the total number of times that the time point was utilized (see Method section). The whole time block was divided into nine sub-temporal regions. The highest spatiotemporal influence index for every three seconds was overlaid on the bar chart (red line). X-axis represents time in seconds. **(A)** Illustrates the temporal influence for overall prediction. Temporal influence in detecting faces and houses are illustrated in **(B)** and **(C)**, respectively.

**Figure 7.**
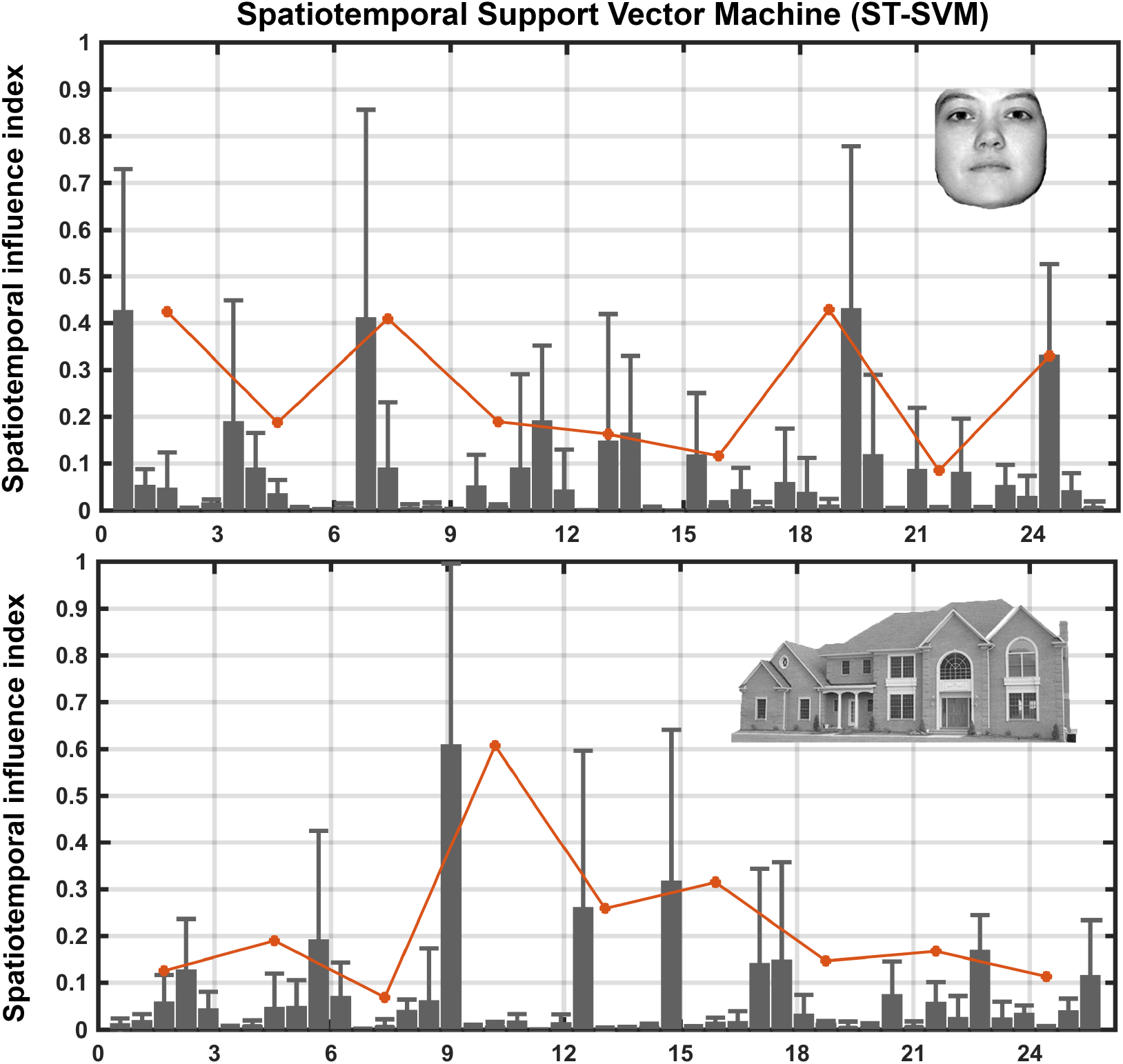
Spatiotemporal influence of each time point using Support Vector Machine (SVM). Bars indicate the mean and standard deviation of spatiotemporal influence index across six runs. Spatiotemporal influence index represents the number of times were each time points was chosen as the most informative time point of the ST combination, normalized over the total number of times that the time point was utilized (see Methods section). X-axis represents time in seconds. Red line indicates the highest spatiotemporal influence index for every three seconds. Temporal influence in detecting faces and houses are illustrated in the top and bottom panels, respectively.

As opposed to ST-RF, in ST-SVM there was no consistent pattern in the performance across runs and categories. In addition, no specific interval with highest accuracy was found. For example, in ST-SVM run 5, excluding the beginning time intervals leads to a noticeable drop in overall performance (i.e. dark region in the top left corner of the map). However, the same time interval led to the highest prediction accuracy in run 6. The aforementioned trials correspond to the interval that starts from the stimuli onset and ends around the peak of HRF (∼5 seconds after stimuli onset).

Comparing Figures 4 and 5, in ST-SVM the highest performance was either equal to or lower than ST-RF. When looking at only the first 10 seconds from the onset (from onset to post HRF peak), ST-RF was superior to ST-SVM. Similar to ST-RF, an increase in prediction accuracy was observed in the later temporal trials in some cases (e.g. after ∼22 seconds, as shown in the right bottom corner of maps).

The results were dominant dependent on the selected ST combination, especially within stimuli categories. For example, in run 2, the prediction accuracy of face stimuli was as high as 90% or as low as 10% depending on the selected ST combination.

### 3.3. TR influence index results

The tri-second maximum TR influence indices were higher in the first quarter of TRs compared to the later TRs (Figure 6). A pattern observed in the tri-second maximum temporal influence across the studied categories using the RF classifier. The interval with highest tri-second maximum influence index contains the peak of the HRF, and the most influential was the 10^th^ TR that is around the peak of HRF, 5.68 seconds. Another peak in the tri-second maximum TR influence index was observed around the 40^th^ TR (∼22 seconds from the stimulus onset), which was consistent across all categories.

In contrast to the tri-second maximum TR influence of ST-RF, there was no specific trend in the tri-second maximum temporal influence across the two categories using ST-SVM. It should be noted that the classification performance for ST-SVM was relatively poor comparing to ST-RF. For the face category, the first TR (0.568 seconds after the stimulus onset) has been selected with a high mean influence level compared to the other TRs across six runs. In the same category, the second best tri-second maximum TR is at the 34^th^ TR (19 seconds after the stimulus onset). The most tri-second maximum influential TR in house category happened after the HRF peak in the 16^th^ TR.

### 3.4. Temporal length in best SpatioTemporal combinations

The top ten accurate ST combinations were from the beginning half intervals (i.e. ∼ the 20^th^ TR) (with the exception of run 5). The ST-RF with highest overall prediction accuracy always contained the peak of HRF across the six runs (the 10^th^ TR). Using ST-SVM, in run 5, an unexpected temporal region appeared to be most informative (around 40^th^ TR). Similarly, using ST-RF in runs 3 and 5 the later temporal domain seemed to be informative.

The most accurate ST-SVMs are no longer than 19 TR and are mainly short in duration across runs (Figure 9). The lengths of ST-RF across the top 1% trials (top ten ST combinations) were longer than ST-SVM on average across cross-validation runs. The measured duration was ∼9s for RF and ∼5s for SVM.

### 3.5. Replicability of the SpatioTemporal results

Consistent with our previous investigation in participant 1, ST resulted in higher prediction accuracy in almost all runs across other participants, compared with single TR technique. The average values of top five prediction accuracies from ST combinations were close to the highest performance, with a small standard deviation (Figure 10). Table 2 showed that the average classification accuracy of ST-RF technique was consistently higher than single TR-RF in four participants across cross-validation runs.

**Table 2.** Average classification accuracy of highest single TR, SpatioTemporal (ST) technique in 4 participants across cross-validation runs. Average overall accuracy of top single TR classification, top five ST technique, using random forest (RF) classifier (rows) across four participants (in columns).

|  | Participant 1 | Participant 2 | Participant 3 | Participant 4 |
| --- | --- | --- | --- | --- |
| Single TR | 69.16 | 63.33 | 67.5 | 62.5 |
| ST | 78.33 | 72.5 | 75 | 75 |

Note that in some cases both single TR and ST settings resulted in poor classification accuracies (e.g. run 2 and 5 of participant 2). Monitoring the skin conductivity (not included here) showed different level of conductance in the aforementioned runs, suggested that the poor performances were most likely due to the decreased engagement of the participant. Note that in some cases the ST combination resulted in a dominant improvement in the prediction accuracy, which was not observed in participant 1. For example, in run 4 of participant 4, ST resulted in 20% improvement of prediction accuracy, compared with 55% prediction accuracy in single TR.

Using RF classifier, the highest overall performance of participant 1, when the first 11 seconds after the stimuli onset was utilized (Figure 10), was similar to when all 25 seconds were used in Figure 2, except for runs 2 and 5. This finding indicates that there is temporal information in between 11 seconds until 25 seconds from the stimulus onset that assist on increasing the MVPC performance.

## 4. Discussion

The effect of ST feature selection on brain decoding was investigated in this study and the obtained prediction accuracies were compared with those obtained using the single TR approach. When considering the single TR scenario, the best decoding performance was achieved using single time point data around the HRF peak. Using ST feature selection, the best sensitivity to each stimulus was 90% or higher that was higher on average than single TR across all cross-validation runs (see ST-RF results in Figure 2B–C). A multi-band EPI pulse sequences was utilized, providing high temporal resolution, which enables rigorous exploration of the temporal domain.

### 4.1. ST features versus single TRs

Results of this study showed that on average, ST feature selection led to higher prediction accuracy compared to the single TR observation. The effect of including the whole ISI (25 seconds) was investigated on the performance of MVPC and the results show that the ten best performing ST combinations do not include the whole ISI (Figures 8 and 9). Furthermore, the highest prediction accuracies were gained with ST combinations that include time points around the peak of the HRF (ST combinations containing from ∼2–11 seconds after the stimuli onset). The discriminative power that was gained from ST combinations within this range was even higher than those combinations containing the entire trial of 25 seconds (used in (Fogelson et al., 2011)). The same conclusion was made when the influence of each time point on the ST combinations was assessed.

**Figure 8.**
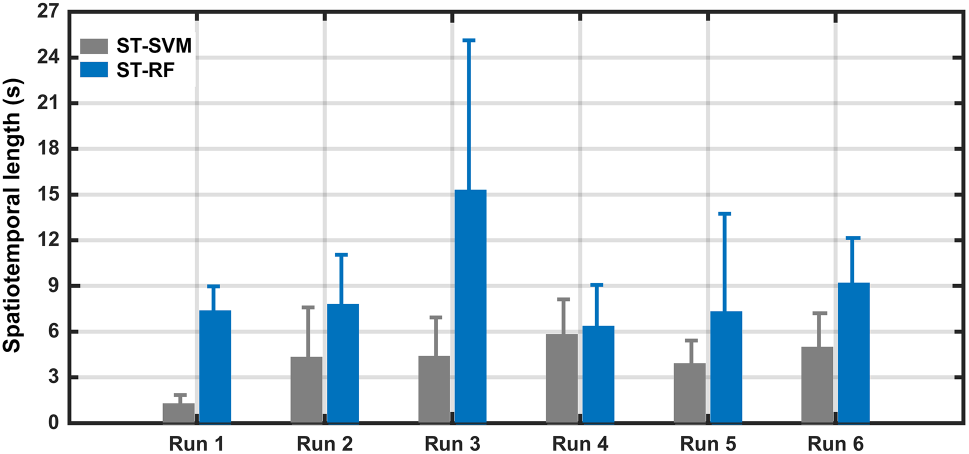
Temporal length of the top ten SpatioTemporal (ST) combinations. Bars indicate the mean and standard deviation of temporal length for ST combinations with highest overall prediction accuracy (top ten ST combinations of each run are considered).

**Figure 9.**
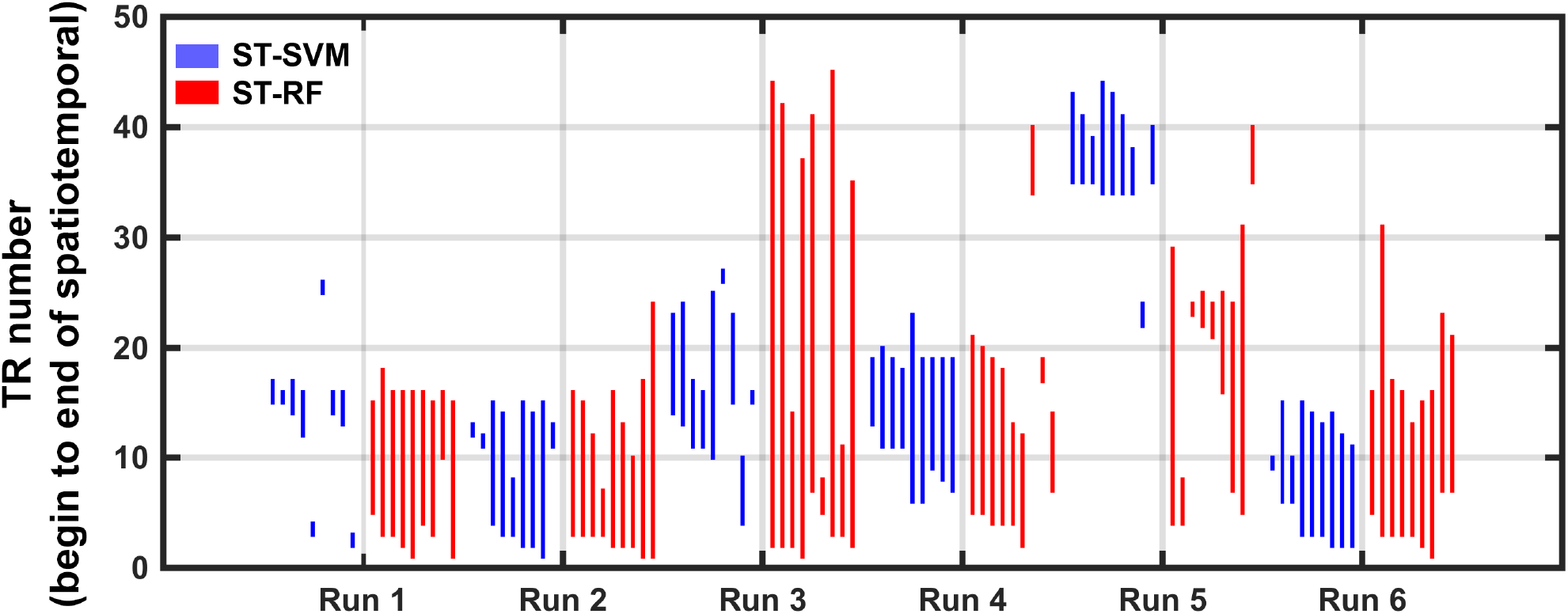
Temporal duration in the top ten SpatioTemporal (ST) trials across six runs. Blue and red lines show the time duration of the ST combination with the top 10 highest prediction accuracy for overall category using SVM and RF, respectively. Each line indicates the start and end point of the ST combination.

In some ST combinations that did not have stimuli specific BOLD activities, the performances of the classifiers were lower than chance. On these cases the classification optimizer fails to diverge, which results to a failure in classification in a systematically biased way. Particularly, when the training data is extremely noisy (in this case, uninformative temporal features) the classification may fit the model to noise and cause a bias.

Findings in this section complied with the strategy used in a recent study where MVPC was employed to decode individual finger movements (Shen et al., 2014). The feature vector was constructed using two successive volumes in the image series for a trial corresponding to the duration close to the peak of the HRF in the studied ROI. Later, the two successive volumes were concatenated to construct spatial-temporal feature vectors (Shen et al., 2014).

### 4.2. Comparison between SVM and RF

Based on the findings in this study, compared to the SVM, RF performs better in MVPC using ST feature selection. RF led to higher prediction accuracy compared to the SVM and showed more consistency across stimuli and runs. No consistent pattern was observed across SVM results from ST combinations. In addition, ST combinations with the highest prediction accuracy from the RF classifier were always longer than those from the SVM (Figure 8-9), suggesting that RF benefits more from temporal information encoded in ST embedding than does the SVM. In general, RF led to higher prediction accuracies across stimuli and runs compared with the SVM, regardless of utilization of ST feature selection.

While SVM algorithms are computationally stable, generalize well, and have been applied successfully to fMRI data (LaConte, Strother, Cherkassky, Anderson, & Hu, 2005), for MVPC with ST feature selection, RF outperformed SVM. The superiority of RF was also reported by Douglas et al. (Douglas et al., 2011) for the conventional case of MVPC analyses. One possible reason for this dominance could be that RF has greater power for handling high-dimensional data compared with SVM. RF holds a unique advantage by employing multiple feature subsets, which is well suited for high-dimensional data. This robustness of RF is largely due to the relative insensitivity of misclassification cost to the bias and variance of the probability estimates in each tree (Hastie, Tibshirani, & Friedman, 2009). In principle, SVMs should be highly resistant to over-fitting but in practice this depends on the careful choice of regularization parameter and the kernel parameters. However, over-fitting can also occur quite easily when tuning the hyper-parameters (Hastie et al., 2009).

### 4.3. HRF peak jittering and temporally averaged BOLD signals

The results of this study were compared with the highest prediction accuracy that was obtained using the single TR around the peak of the HRF. A temporal range was considered around the peak of the HRF, rather than the canonical peak. Time point by time point, MVPC showed that the peak of classification accuracy is around the peak of the region-average HRF (Kohler et al., 2013); in some regions prior to and in some regions after the region-average HRF peak. By performing the comparison against the highest prediction accuracy around the peak (based on the findings of Kohler et al. (Kohler et al., 2013)), ST-based results are compared against those from the state-of-the-art approaches. It would be interesting to investigate the hypotheses in this experiment applied to other brain regions.

### 4.4. High decoding accuracy at the end of ISI

The provided classification weight vectors from ST-based input data identifies when class-discriminating information arises, indicated by the TR influence index (Figure 6). Using RF, the tri-second average TR influence index in the first part of the trials in the IT conformed to the temporal pattern in the canonical double gamma HRF. In addition to the time around the peak of HRF, high temporal influence was observed around 23 seconds after stimuli onset. The time at which this observation occurred is around the time when negative undershoot is almost passed according to the canonical double gamma HRF (Friston et al., 1994). The participant’s task in this study was to perform a one back repetition detection task from trial to trial. For each trial the participant attempts to retain in their memory details of the stimuli introduced in that trial, and then waits for the next trial in which they perform the one back repetition detection task. Category expectation was found to be affecting the baseline and stimulus evoked activity in IT (Puri, Wojciulik, & Ranganath, 2009). Expectation was reported to be related to a degree of certainty about an upcoming stimulus (Cisek & Kalaska, 2010), meaning that if a certain stimulus category has a high probability of appearance, preparatory processes of expectation can facilitate its detection and the associated responses (Cisek & Kalaska, 2010). It was reported that the baseline activity level in subcortical regions in IT (FFA and PPA) was higher during expectation of the preferred (e.g. face for FFA) versus non-preferred category (Egner, Monti, & Summerfield, 2010; Herwig, Abler, Walter, & Erk, 2007; Puri et al., 2009). Therefore, a high chance of correspondence exists between the high discrimination power in the 23^rd^ second of the trial, and the expectation mechanism in IT. Observing such an effect in the TR influence highly depends on the experimental design, brain region, the task, and the inter stimulus interval. Note that the influence of the TRs suspected to be related to expectation was much lower than the TRs around the peak of the HRF. The latter finding invites further investigations on the effect of category expectation using MVPC.

### 4.5. Improved performance across participants

In this study, a rigorous investigation was performed on the effect of ST feature selection on the MVPC performance for one of the participants. Later, the analysis conclusion was validated on the rest of the participants. Participant 1 was chosen as the subject for a detailed comparison as the MVPC performance in single TR for participant 1 was overall higher than other subjects of this study. For the other participants, only the first 11 seconds of each trial were utilized to investigate the effect of ST feature selection. This time interval was also employed for stimuli design in a previous study (Kohler et al., 2013), and our initial investigations recommended that this temporal range embraces highly decodable dynamics. All other 990 ST combinations were derived for each of the 4 participants. Of course, higher prediction accuracy than what is reported (Figure 10) might have been obtained if the entire temporal domain was considered (e.g. comparing ST-RF results in Figure 1A and Figure 10A). But here, our aim was to show that even within this range an improvement in prediction accuracy could be obtained by the use of ST feature selection.

**Figure 10.**
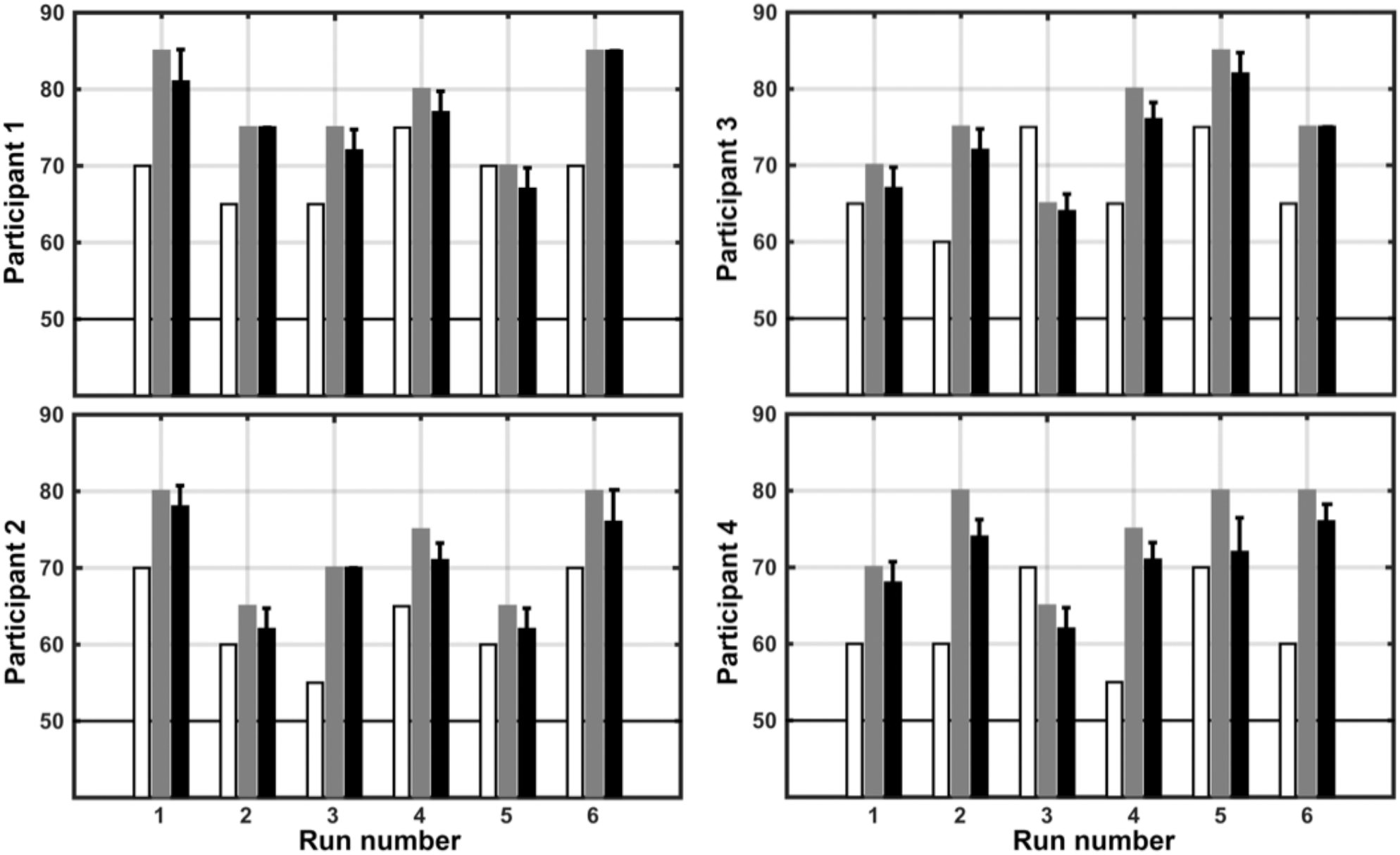
Highest classification accuracy of single TR versus SpatioTemporal (ST) technique across 4 participants. Overall accuracy of single TR classification is compared with ST technique using random forest (RF) classifier (RF-ST). White bar demonstrates the overall performance of the single TR classification. Grey bar demonstrates the highest classification accuracy using RF-ST technique considering only at the first 11 seconds from the stimuli onset. Black bar shows the mean and standard deviation of top five prediction accuracies within the aforementioned temporal domain.

### 4.6. Implications of temporal feature selection on brain decoding

Brain decoding not only allows for combinational effects across voxels, but also has applications in brain-computer interfacing (Davis & Poldrack, 2013; Van De Ville & Lee, 2012). However, due to the challenges related to sensitivity and specificity, clinical justification has not fully been achieved yet. But an improved brain decoding technique can hasten the transition time of MVPC from laboratory to clinic.

The findings of this study open the way for further investigations into the understanding of category learning dynamics, where the stimuli space will change over time as a result of learning (Davis & Poldrack, 2013). A potential application of incorporating our ST approach into MVPC would be to see if by embedding the dynamic patterns, the time at which the brain starts shaping a motor decision and unconscious mental processes could be decoded in a shorter temporal duration (Soon, Brass, Heinze, & Haynes, 2008). Our observations have methodological implications for selecting the time at which to perform classification analyses. Throughout IT there can be systematic differences in the temporal dynamics of classification accuracy that could be investigated by ST embedding. However, it remains to be seen as to whether these findings will generalize to other areas and other stimuli.

One of the limitations of MVPC studies is the reported weak correlation of inter-subject and intra-subject measured fMRI responses to the same stimuli (Chen et al., 2014). In addition, it is challenging to establish a correspondence between selected voxels/ROIs across participants in a dataset. A challenge in this study was to tax the subject’s attention considering our long ISI. It is desirable to create an experiment design that required the subjects to spend less time inside the scanner and to watch stimuli for as long as time allows (Franklin, 2012). However, the aim of this study was to investigate a wide range of temporal information, which can lead to such ISI design.

While improvement was gained on average by utilizing ST feature selection, no specific combination consistently outperformed others across runs and participants. This provides a motivation for the future direction of our investigation on brain decoding using ST feature selection, and that is to find a systematic optimum way to extract ST features.

In conclusion, on average the ST feature selection led to higher classification performance compared with single TR observation. This study explored the temporal domain and found that the discriminative power increased when time points around the peak of the HRF were included in the ST combination. In particular, ST combinations taken from between ∼2–11 seconds after the stimuli onset outperformed the rest of the ST combinations (including those combinations containing the entire trial, i.e. the whole trial embedded ST). Our assessment of the importance of time points in ST combinations confirmed the latter conclusion about the most important time points. Based on the evaluation criteria of this study (MVPC prediction performance and the conforming temporal pattern in TR influence index with canonical double gamma HRF), the findings in this study suggest RF as the classifier of choice over SVM for brain decoding using ST feature selection. RF led to higher prediction accuracy, and showed more consistency across stimuli and runs. RF also benefits more from ST information compared with SVM (the length of the highest performance ST combination was always longer with RF than with SVM).

## Acknowledgement

This work was support by W.M. Keck Foundation, National Institute of Health (5T90DA022768), and by Staglin Center for Center for Cognitive Neuroscience. Choupan J. was supported by University of Queensland International PhD Scholarship.

